# Respiratory tract *Moraxella catarrhalis* and *Klebsiella pneumoniae* can promote pathogenicity of myelin-reactive Th17 cells

**DOI:** 10.1101/2023.03.30.534905

**Authors:** Jenny M. Mannion, Benjamin M. Segal, Rachel M. McLoughlin, Stephen J. Lalor

## Abstract

The respiratory tract is home to a diverse microbial community whose influence on local and systemic immune responses is only beginning to be appreciated. The airways have been linked with trafficking of myelin-specific T cells in the pre-clinical stages of experimental autoimmune encephalomyelitis (EAE), an animal model of multiple sclerosis. Th17 cells are important pathogenic effectors in MS and EAE but are innocuous immediately following differentiation. Upregulation of the cytokine GM-CSF appears to be a critical step in their acquisition of pathogenic potential, but little is known about the mechanisms that mediate this process. Here, primed myelin-specific Th17 cells were transferred to congenic recipient mice prior to exposure to various human respiratory tract-associated bacteria and T cell trafficking, phenotype and the severity of resulting EAE monitored. Disease was exacerbated in mice exposed to the Proteobacteria *Moraxella catarrhalis* and *Klebsiella pneumoniae,* but not the Firmicute *Veillonella parvula*, and this was associated with a significant increase in GM-CSF^+^ and GM-CSF^+^IFNγ^+^ ex-Th17-like donor CD4 T cells in the lungs and CNS of these mice. These findings support the concept that respiratory bacteria may contribute to the pathophysiology of CNS autoimmunity by modulating pathogenicity in crucial T cell subsets that orchestrate neuroinflammation.

## Introduction

The respiratory tract is home to a diverse microbial community that is dominated by species from the phyla Bacteroidetes, Proteobacteria, Firmicutes, Actinobacteria and Fusobacteria [1]. The composition of the airway microbiota varies from site to site because of the diverse physiological conditions that exist along the respiratory tract. In the past decade, associations have been made between the composition of the airway microbiota and the pathophysiology of chronic respiratory diseases including asthma [2], chronic obstructive pulmonary disease (COPD) [3] and cystic fibrosis (CF) [4]. In addition, the lung microbiota is implicated in the early development of the systemic inflammatory disease rheumatoid arthritis (RA) [5]. Beyond association, little is known about the immunomodulatory capacity of discrete members of the respiratory microbiota, and research is now shifting to elucidation of the mechanisms by which individual strains of bacteria might regulate disease susceptibility and immunopathology.

A growing literature has linked the lungs with the pathogenicity of immune cells that orchestrate CNS autoimmunity [6–9]. Multiple sclerosis (MS) is a chronic inflammatory disease of the central nervous system (CNS) widely believed to be instigated by the infiltration of autoreactive T cells and other immune cell subsets into the CNS which attack the myelin sheath surrounding nerve axons, resulting in the formation of an inflammatory plaque and reduced signal conductance [10, 11]. High rates of discordance amongst monozygotic twins indicate that environmental factors play a crucial role in promoting the development of MS, and epidemiological studies have long associated relapses in MS patients with systemic infection. [12–14]. Studies using germ-free mice in experimental autoimmune encephalomyelitis (EAE), an animal model that recapitulates many of the clinical features of MS, have also implicated microbial factors as important environmental stimuli in promoting CNS autoimmune inflammation [15, 16]. How specific members of the airway microbiome might promote pathogenicity in MS and EAE is unknown.

Recent studies have demonstrated that the lung microenvironment can promote a pathogenic phenotype in CD4^+^ T cells as they traffic to the CNS. In a rat model of EAE, it was demonstrated that the gene expression profile of activated myelin-reactive T cells is altered during their migration through the lungs to the CNS, promoting the ability of those cells to cross the blood-brain barrier and mediate neuroinflammation [6]. Separately, it was found that expansion of a pro-inflammatory population of granulocytic myeloid-derived suppressor cells (MDSC) in the lungs of mice during the development of EAE enhanced the pathogenicity of a key population of IL-17-expressing myelin-specific T (Th17) cells [9]. Indeed, myelin-reactive Th17 cells have emerged as important pathogenic effector cells in both MS and EAE. IL-17-producing CD4^+^ T cells accumulate at increased frequencies in the blood and cerebrospinal fluid (CSF) during MS, particularly during relapses [17–19]. In addition, messenger RNA encoding IL-17 is upregulated in the CNS of individuals with MS, and IL-17^+^ CD4^+^ T cells have been detected in MS lesions by immunohistochemical analysis [20, 21]. Th17 cells are induced in the presence of IL-1, IL-6 and TGFβ, express the transcription factor RORγt and secrete the cytokines IL-17A, IL-17F, IL-21 and IL-22, but are innocuous immediately following differentiation [22]. IL-23 signaling is critical for Th17 cell pathogenicity and IL-23-driven conversion to a GM-CSF and IFNγ producing “ex-Th17” cell phenotype appears to be a critical step in their acquisition of pathogenic potential [23–25]. During EAE, GM-CSF promotes the development of an inflammatory phagocyte phenotype in monocyte-derived effector cells and, like IL-23-deficient mice, mice lacking GM-CSF are relatively resistant to EAE [26–28]. Naïve CD4 T cells do not express IL-23R, however, and only upregulate the receptor following expression of the major Th17 cell transcription factor RORγt [22, 29]. When and where these cells are exposed to this pathogenic signal is unknown.

In the current study, careful kinetics analyses revealed that myelin-specific Th17 cells rapidly migrate to the lungs following transfer to naïve mice. In support of a role for commensal microbes in conferring encephalitogenicity, Th17 cells induced attenuated demyelinating disease upon transfer to germ-free mice, compared to that observed in specific pathogen-free (SPF) hosts. Importantly, we demonstrate that airway exposure to the Proteobacteria *Moraxella catarrhalis* and *Klebsiella pneumoniae* significantly exacerbates clinical disease in mice with EAE. Aggravated EAE was associated with increased frequency of donor CD4 T cells expressing GM-CSF or IFNγ, or both cytokines together, in the lungs during the preclinical stages of disease. Employing *in vitro* Th17 cell re-activation assays, we show that this switch to an “ex-Th17” cell phenotype can be directly promoted by exposure of antigen-presenting dendritic cells (DC) to these discrete bacterial strains. Together these findings support the hypothesis that perturbations in the composition of the respiratory tract microbiota regulates acquisition of pathogenicity by myelin-reactive Th17 cells and the development of CNS autoimmune inflammation.

## Results and Discussion

### Donor Th17 cells traffic in the lungs prior to their accumulation in the CNS and their pathogenicity is modulated by the commensal microbiota

Following transfer of *in vitro* polarized MOG-specific Th17 cells to naïve syngeneic mice, there is a period of approximately one week before onset of clinical signs of EAE. The reasons for this delay are unknown and unexpected given the activated state of the autoantigen-specific T cells that are transferred. Hence, we assessed trafficking and accumulation of transferred cells in the CNS and peripheral tissues daily between the time of adoptive transfer and the development of clinical EAE. MOG-specific Th17 cells were generated *in vitro* and 1×10^6^ enriched CD45.1^+^ donor CD4^+^ T cells injected i.p. to naïve congenic CD45.2^+^ C57BL/6 hosts. The majority of mice displayed initial signs of disease 7d post-transfer, approximately 48h following initial infiltration of donor CD4 T cells into the brain (mean=11934 donor CD4 T cells/brain) and spinal cord (mean=387 donor CD4 T cells/SC) on d5 post-transfer (Fig. 1A). Examination of mucosal and peripheral lymphoid tissues during the pre-clinical stages of EAE revealed significant accumulation of donor CD4 T cells in the lungs and mesenteric lymph nodes between d3 and d6 following transfer, but not in the Peyer’s patches or the cervical or inguinal lymph nodes (Fig. 1B).

**Figure 1.**
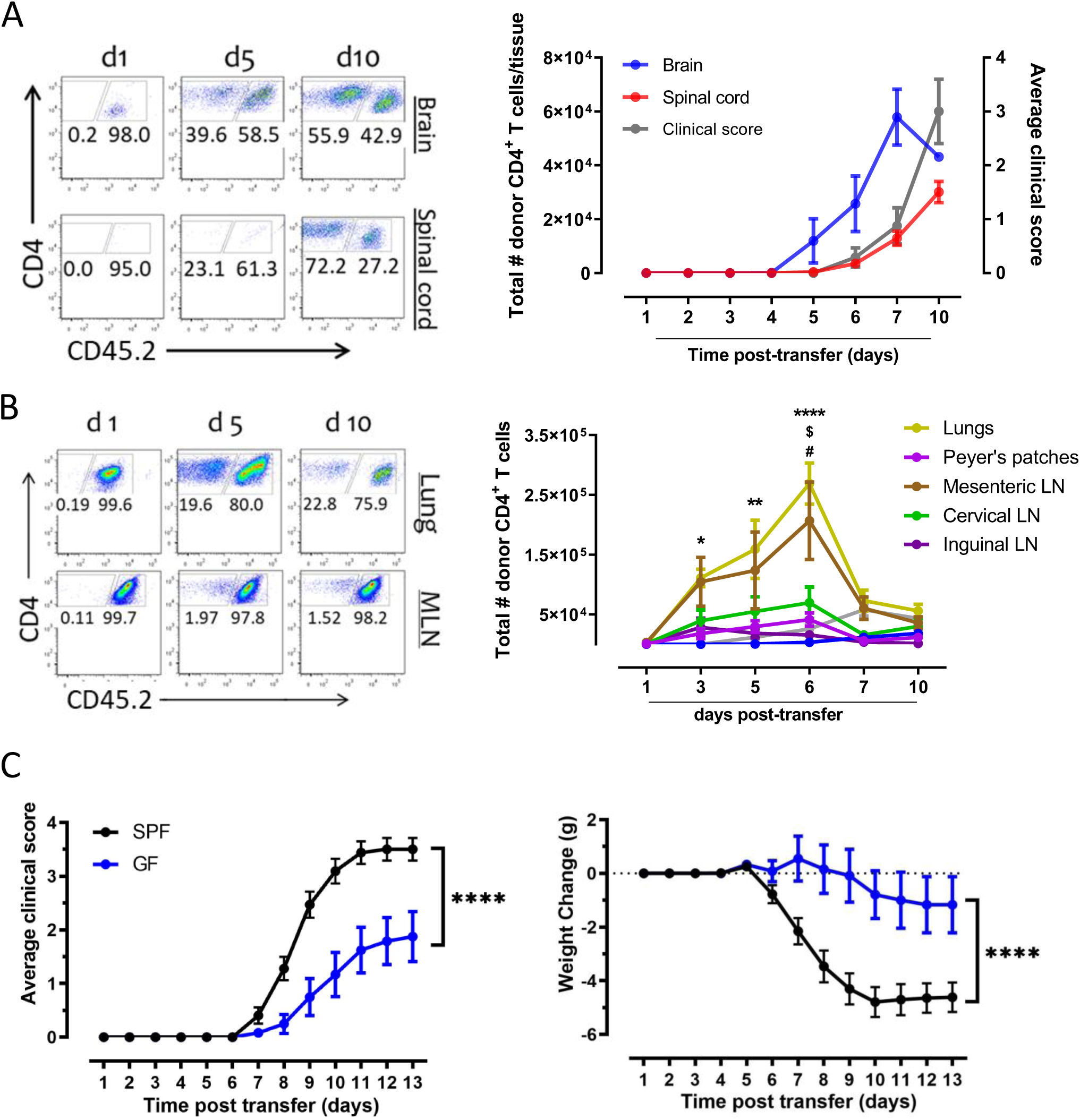
Myelin-reactive Th17 cells traffic through mucosal tissues in the preclinical stages of EAE and are less encephalitogenic when transferred to germ-free mice. Th17-polarized MOG-specific T cells (from CD45.1 donor mice) were adoptively transferred (1X10^6^ CD4^+^ T cells/mouse) to naïve congenic (CD45.2) SPF recipients via i.p. injection and animals weighed daily and monitored for clinical signs of disease. At the indicated time points, single cell suspensions were prepared from brains, spinal cords, lungs (*), Peyer’s patches (^$^), mesenteric (^#^), cervical and inguinal lymph nodes from individual mice and stained with fluorochrome-conjugated antibodies specific for CD4, CD45.1 and CD45.2. Representative FACS plots and total number of donor CD4 T cells per brain and spinal cord (A) or lung and mesenteric lymph nodes (MLN) cells (B). Numbers are % of CD4^+^ gate and total viable congenic donor CD4 T cells/tissue (A & B: n=4-12/group from 3 independent experiments). Statistical analysis comparing donor CD4 T cells/tissue over time was performed using one-way ANOVA with multiple comparisons. * P<0.05, ** P<0.01, **** P ≤ 0.0001 vs d1 post-transfer. Th17-polarized MOG-specific T cells were adoptively transferred (1X10^6^ CD4^+^ T cells/mouse) to naïve SPF or germ-free (GF) recipients via i.p injection and animals weighed daily and monitored for clinical signs of disease (C; n=12-16/group from 4 independent experiments). Statistical analysis comparing clinical scores and mouse weight over time was performed using two-way ANOVA with multiple comparisons. **** P ≤ 0.0001 vs SPF recipients.

Our observation that donor Th17 cells preferentially accumulate at mucosal sites in the pre-clinical stages of EAE supports the hypothesis that microbial factors may influence Th17 cell pathogenesis and disease development. To assess the impact of the commensal microbiota specifically on the effector function of Th17 cells, we transferred 1×10^6^ purified Th17-polarized CD4 T cells to syngeneic germ-free mice and monitored them daily for clinical signs of disease.

We found that germ-free recipients of SPF-derived Th17 cells exhibited significantly reduced incidence and severity of disease compared to SPF recipients of the same Th17 cells (Fig. 1C). Attenuated EAE in germ-free hosts was also associated with significantly reduced weight-loss (Fig. 1C), an objective measure of disease severity and sickness behavior in mice experiencing an acute inflammatory challenge [30]. Together, this data indicates that myelin-specific Th17 cells may be exposed to commensal microbes at mucosal sites, including the lungs, that can influence their accumulation and/or pathogenicity in the CNS (Fig. 1B) [6, 9]. Unlike the attenuated disease observed in actively induced or spontaneous models of EAE in germ-free mice that exhibit defective immune development, this study also indicates that commensal microbes can specifically regulate the effector function of myelin-specific Th17 cells [15, 16].

### IL-23 expression is strongly induced *in vitro* and *in vivo* by species from the phyla Bacteroidetes, Proteobacteria and Fusobacteria

Overall, little is known about the immunomodulatory capacity of individual members of the airway microbiota. In EAE, however, IL-23 signalling is critical for the conversion of Th17 cells to a GM-CSF and IFNγ-expressing pathogenic ex-Th17 cell phenotype [22–25, 31]. Hence, to identify bacteria that might impact Th17 cell pathogenicity in our model of EAE, we investigated the capacity of 29 individual strains of bacteria that commonly colonize the human respiratory tract (Supp. Table 1) to stimulate IL-23 secretion by antigen-presenting cells that could play a role in Th17 cell activation and plasticity. We found that stimulation of bone marrow-derived DC (BMDC) with live stains of Bacteroidetes, Proteobacteria and Fusobacteria *in vitro* strongly promotes IL-23 production (Fig. 2A). None of the phyla Firmicutes or Actinobacteria tested in this study exhibited this ability, although *Streptococcus mitis* and *Streptococcus pneumoniae* could stimulate IL-12p70 expression by DC. Interestingly, the strong IL-23-inducing strains *Neisseria cinerea* (CCUG346G), *K.pneumoniae* (ATCC43816) and *Fusobacterium nucleatum* (ATCC25586) did not induce IL-12p70 expression in these assay conditions. All bacteria stimulated strong TNFα responses, demonstrating their immunogenic potential, with limited cytotoxicity (Supp. Fig. 1B). We failed to detect IL-23 or IL-12p70 expression by bone marrow-derive macrophages (BMM; Supp. Fig. 1A) or FACS-sorted alveolar macrophages (AM; Supp. Fig. 1D-E) from the lungs of naïve C57BL/6 mice in response to any of the bacterial strains tested, despite strong TNFα production and low cytotoxicity (Supp. Fig. 1C). This indicates that DC may be a more likely source of IL-23 following exposure to these human respiratory bacteria, and is in line with previous data indicating a lack of IL-23 expression by murine AM [32]. Human lung macrophages may respond differently, however, given their repeated exposure to air pollutants and respiratory infections from early life.

**Figure 2.**
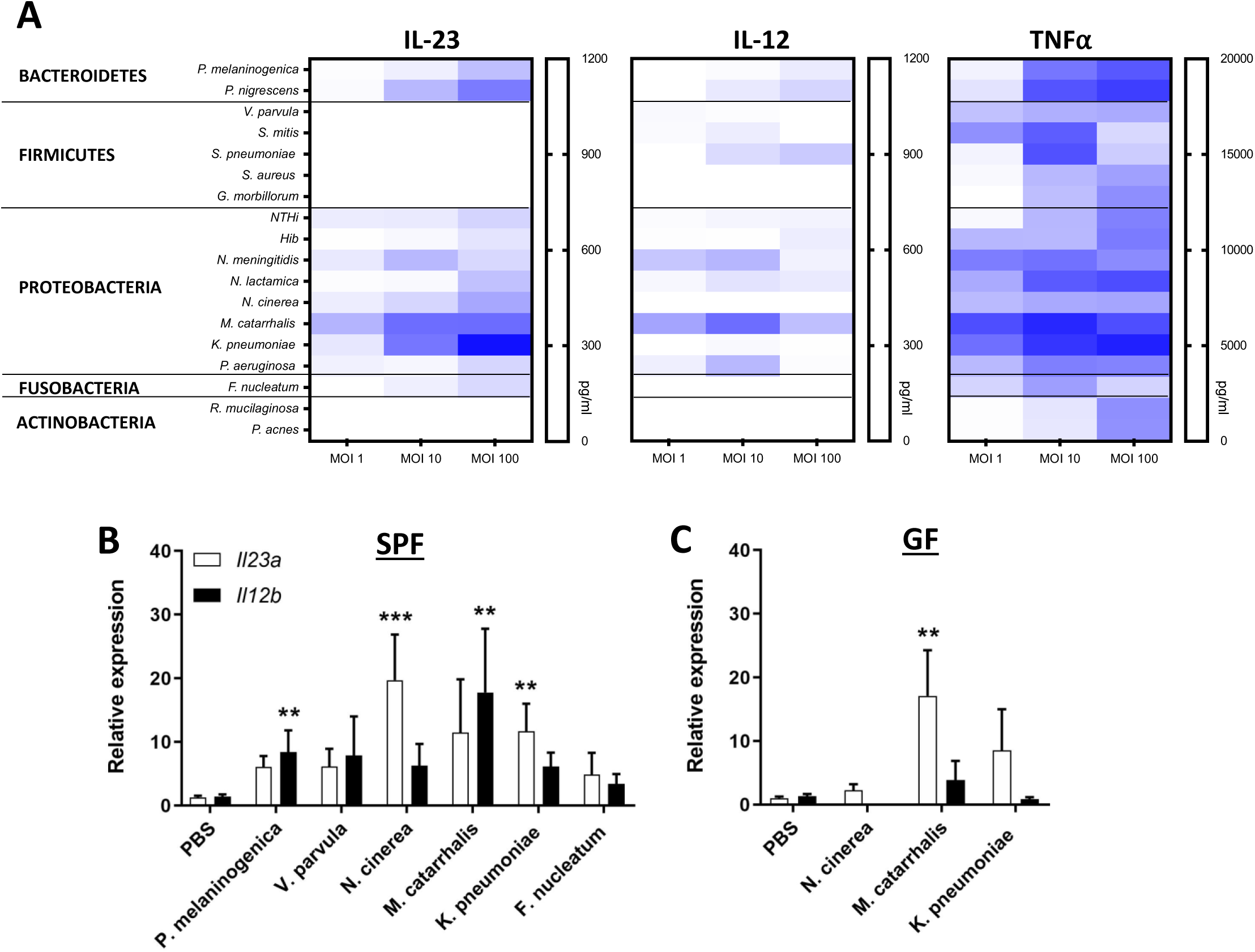
Bacteroidetes, Proteobacteria and Fusobacteria species promote IL-23 expression by dendritic cells *in vitro* and *in vivo* in the lungs of exposed mice. BMDC were seeded in triplicate at 2X10^5^ cells/well in a 96 well plate and rested for 3 h before exposure to indicated bacteria at MOI 1, 10 and 100. Cells were treated with gentamicin (100μg/ml) after 1h and supernatants collected 23h later. The concentration of IL-23, IL-12p70 and TNFα was quantified by ELISA. Results are presented as a heat map indicating average cytokine concentration ml^−1^ (A; n=3-9/group). Groups of SPF (B) and GF (C) mice were exposed to the indicated bacteria via i.n. administration of 20μl PBS containing 1×10^6^ CFU, or PBS alone. 24h post-administration, RNA was extracted from lung tissue and gene expression for *Il23a* and *Il12b* assessed by RT-PCR. mRNA values are expressed as mean fold change ± SEM, compared to PBS-administered controls, after normalising to the housekeeping gene 18S rRNA (n=4-8/group from 2-3 independent experiments). Statistical significance was determined by Student’s *t* test. ** P<0.01, *** P<0.001 vs PBS administered controls.

To test if the observed cytokine responses are replicated *in vivo*, we exposed mice to selected bacteria via the intranasal (i.n.) route and found significantly increased *Il23a* and *Il12b* gene expression in the lungs of mice exposed to the Bacteroidetes species *Prevotella melaninogenica* and the Proteobacteria *N.cinerea, M.catarrhalis* and *K.pneumoniae*, compared to PBS-administered controls (Fig. 2B). Similarly, *M.catarrhalis* and *K.pneumoniae* directly promoted increased *Il23a* gene expression in the lungs of germ-free mice that lack a complex microbiota which could otherwise modulate observed immune responses (Fig. 2C).

### Airway exposure of Th17 cell recipient mice to discrete respiratory-associated Proteobacteria exacerbates clinical signs of EAE

To determine how exposure to individual strains of IL-23-stimulating human respiratory bacteria might modulate the effector function of Th17 cells in the passive transfer model of EAE, we transferred 1×10^6^ *in vitro* polarised MOG-specific Th17 cells to SPF mice and, 2d later when these cells begin to accumulate in the lungs (Fig. 1B), exposed different groups to selected bacteria via the i.n. route. Exposure to *V.parvula*, typically associated with the healthy human airways and which did not induce any IL-23 secretion in innate immune cells *in vitro* (Fig. 2), had no effect on the clinical course of disease or lead to any form of sickness behaviour in mice with EAE (Fig. 3A). However, airway exposure of Th17 cell recipient mice to *M.catarrhalis* (Fig. 3B) or *K.pneumoniae* (Fig. 3C) led to a significant increase in EAE severity, associated with earlier onset and more severe clinical signs of disease than PBS-administered controls. Mice exposed to these bacteria also lost significantly more weight than their PBS-administered counterparts, indicating both neurological and behavioural impacts following acute immune challenge with these respiratory bacteria. Moreover, transfer of myelin-specific Th17 cells to germ-free mice monocolonised with *K.pneumoniae* from birth resulted in increased incidence (77% versus 55%) and severity of EAE compared to germ-free recipients of the same Th17 cells (Fig. 3D). Importantly, the observed deficits were not simply a result of the inflammatory response or sickness behaviour caused by the introduction of a bacterial species into the airways because naïve mice administered either *M.catarrhalis* (Fig. 3B) or *K.pneumoniae* (Fig. 3C) alone did not experience any weight changes or discernible adverse effects in response to the bacteria. This data supports the concept that long-term colonisation or recurrent exposure to discrete strains of human respiratory bacteria can enhance the pathogenicity of myelin-specific Th17 cells and exacerbate clinical signs of EAE.

**Figure 3.**
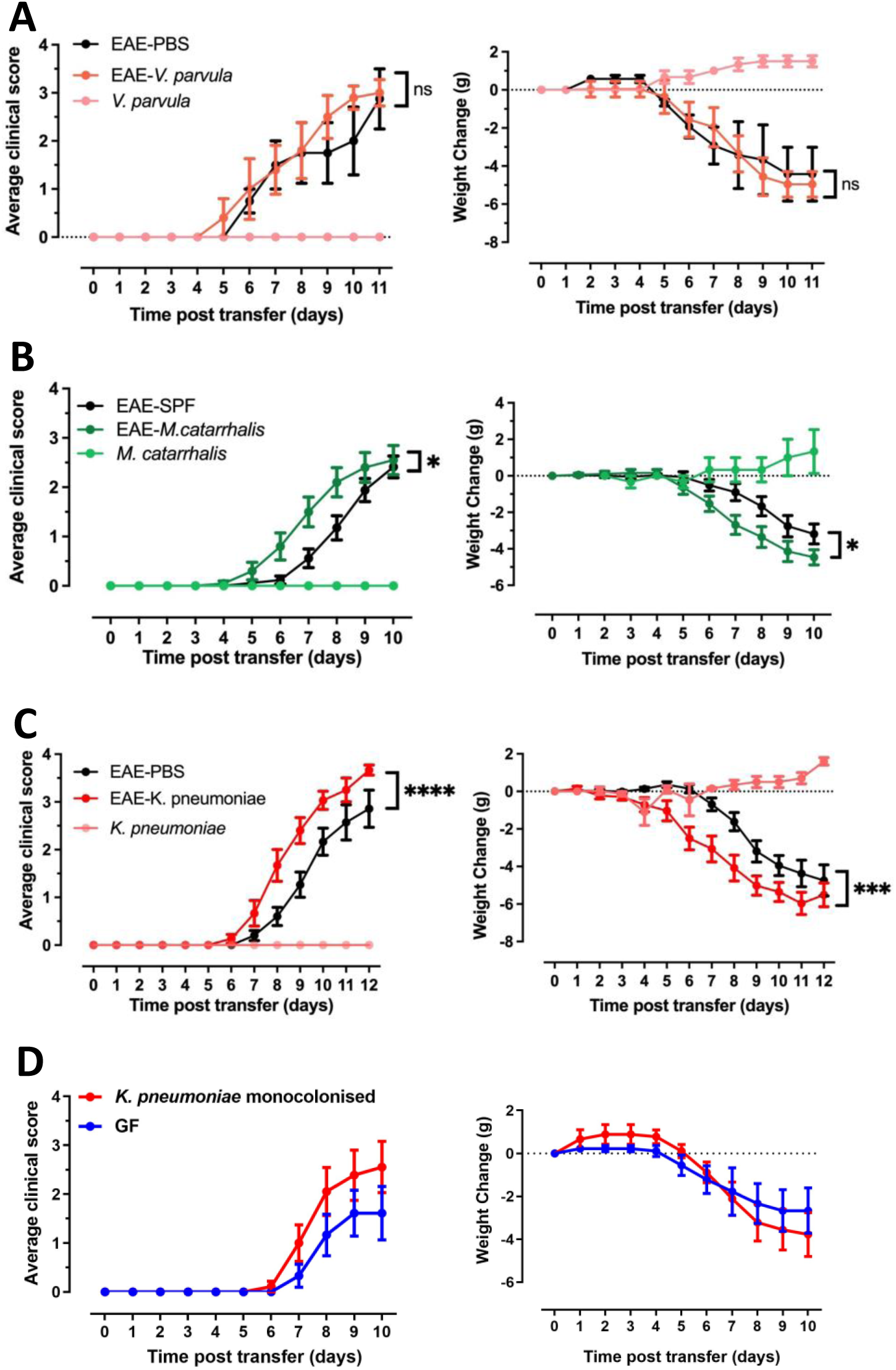
Airway exposure to discrete Proteobacteria species exacerbates Th17 cell-mediated EAE. Th17-polarized MOG-specific T cells were adoptively transferred (1X10^6^ CD4^+^ T cells/mouse) to naïve SPF recipient mice via i.p. injection. 2d after transfer, groups of recipient mice were exposed to the indicated bacteria via i.n. administration of 20μl PBS containing each bacterium, or PBS alone. Mice were weighed daily and monitored for clinical signs of disease. Results are mean clinical score or mouse weight +/− SEM (A-C; n= 5-18 mice/group from 2-5 independent experiments). Th17-polarized MOG-specific T cells were adoptively transferred (1X10^6^ CD4^+^ T cells/mouse) via i.p. injection to naïve GF mice or mice monocolonised with *K.pneumoniae* from birth. Results are mean score or mouse weight +/− SEM (D; n=9 mice/group from 2 independent experiments). Statistical analysis comparing clinical scores and mouse weight over time was performed using two-way ANOVA with multiple comparisons. * P<0.05, *** P<0.001, **** P ≤ 0.0001 vs germ-free recipients.

### Airway exposure to *M.catarrhalis* and *K.pneumoniae* is associated with increased expression of pathogenic cytokines by donor CD4 T cells in the spinal cord and lungs of recipient mice

Th17 cells have been reported to undergo conversion from an IL-17-dominant cytokine phenotype to an IFNγ- and GM-CSF-producing phenotype during the pathogenesis of EAE. In fact, the majority of IFNγ-expressing CD4 T cells in the CNS at the height of disease originally developed along the Th17 lineage and converted to an IFNγ^+^ phenotype following exposure to the cytokine IL-23, while it has since emerged that GM-CSF expression by myelin-specific CD4 T cells promotes disease development [23, 28, 31]. We found that airway exposure of SPF recipients of MOG-specific Th17 cells to *K.pneumoniae* was associated with significantly greater accumulation of IL-17^+^, IFNγ^+^ and GM-CSF^+^ CD4 T cells in the spinal cord at peak disease (Fig. 4A). Tracking congenic donor CD4 T cells revealed that *M.catarrhalis* exposure led to significantly increased frequency of GM-CSF^+^ and IFNγ^+^GM-CSF^+^ double-positive donor CD4 T cells in the spinal cord at the peak of EAE, compared to PBS-administered recipients of the same donor Th17 cells (Fig. 4B).

**Figure 4.**
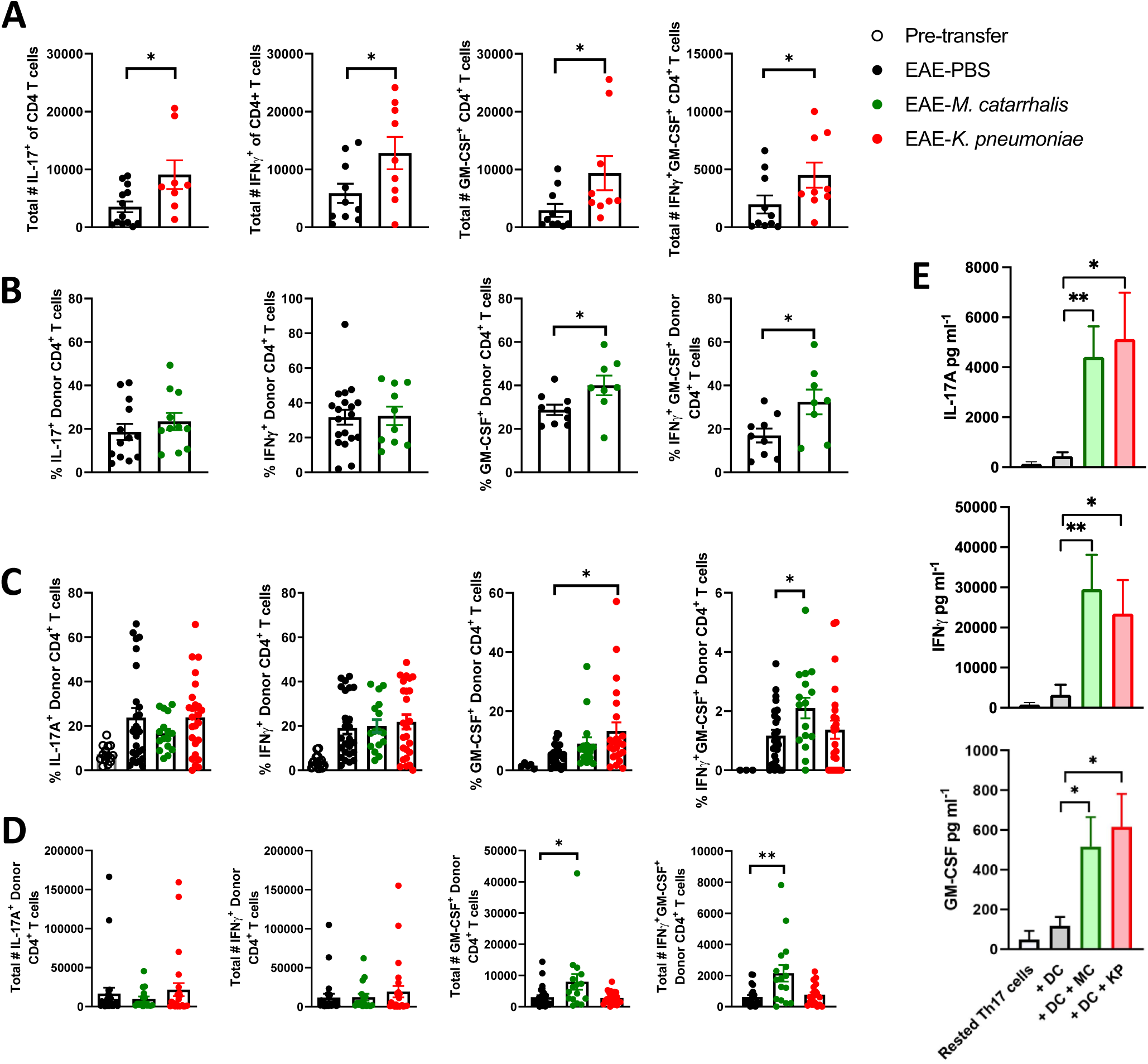
Exposure to *K.pneumoniae* and *M.catarrhalis* promotes expression of the pathogenic cytokines GM-CSF and IFNγ by donor CD4 T cells in the lungs and spinal cord of recipient mice. CD45.1^+^ donor Th17 cells were adoptively transferred (1X10^6^ CD4^+^ T cells/mouse) to congenic CD45.2 SPF hosts via i.p. injection. 2d after transfer, groups of T cell recipient mice were exposed to bacteria via i.n. administration of *M.catarrhalis* or *K.pnumoniae*, or PBS alone. 6d post-transfer (pre-clinical) and at peak disease (d10), mice were perfused intracardially and CNS and lung tissues harvested. Single cell suspensions were stimulated with PMA and ionomycin with brefeldin A for 4 h, stained for surface CD3, CD4, CD45.1 and intracellular IL-17A, IFNγ and GM-CSF, and analysed by flow cytometry. Results are total number (A & C) and mean frequency (B & D) of IL-17A^+^, IFNγ^+^, GM-CSF^+^ and IFNγ^+^GM-CSF^+^ double positive CD4^+^ T cells in the spinal cord (d10; A & B) and lungs (d6; C & D) of Th17-recipient mice exposed to *M.catarrhalis* or *K.pnumoniae*, respectively. Results are expressed as mean +/− SEM (n = 9-25/group from 3-5 independent experiments). Magnetically enriched MOG-specific Th17 cells were rested for 72h following polarization. BMDC were seeded in triplicate at 2X10^5^ cells/well in a 96 well plate and rested for 3h before being pulsed with MOG_35-55_ and exposed to *M.catarrhalis* or *K.pneumoniae* at MOI 100, or media alone. BMDC were treated with gentamicin (100 μg/ml) after 1 h and re-cultured with media containing gentamicin and 2X10^5^ rested Th17 cells for a further 24h. Supernatants were collected and the concentration of IL-17A, IFNγ, and GM-CSF quantified by ELISA (E). Results are expressed as mean ± SEM (n=8/group from 8 independent Th17 polarisation experiments). Statistical significance was determined using Student’s *t* test. * P<0.05, ** P<0.01 vs media only controls.

Investigating development of this pathogenic phenotype by CD4 T cells in the lungs, prior to their infiltration of the CNS, we found that, while all donor cells that had accumulated in the lungs had upregulated GM-CSF and IFNγ expression relative to that expressed at the time of transfer, exposure to *K.pneumoniae* and *M.catarrhalis* enhanced the frequency (Fig. 4C) and, in the case of *M.catarrhalis*, total number (Fig. 4D) of GM-CSF^+^ and GM-CSF^+^IFNγ^+^ double-positive donor CD4^+^ T cells in the lungs compared to PBS-administered controls, 6 d following transfer. Strikingly, host CD4^+^ T cells were unresponsive to bacterial stimulation in terms of IL-17A, IFNγ and GM-CSF production (Supp. Fig. 2A). Moreover, we did not detect any changes in cytokine expression by donor CD4 T cells that accumulated in the spleens of mice exposed to *M.catarrhalis* or *K.pneumoniae* in the preclinical stages of disease (Supp. Fig. 2B). To test whether DC stimulation by these respiratory bacteria can directly promote conversion to this pathogenic “ex-Th17” phenotype, we established a co-culture assay system whereby myelin-specific Th17 cells generated *in vitro* were rested for 3 d before re-culture with BMDC that had been exposed to *M.catarrhalis* or *K.pneumoniae*, or BMDC alone. Using this system, we show that bacteria-stimulated BMDC drive a significant increase in secretion of GM-CSF, IFNγ and IL-17A by myelin-specific Th17 cells, compared to cells reactivated by BMDC alone (Fig. 4E). Together, this data suggests that *M.catarrhalis* and *K.pneumoniae* can modulate the cytokine phenotype and encephalitogenicity of primed CD4 T cells as they traffic through the lungs.

Our results indicate that perturbations in the airway microbiota can exacerbate disease in a preclinical model of MS. We demonstrate that airway exposure to *M.catarrhalis* and *K.pneumoniae* promotes the upregulation of GM-CSF and IFNγ in myelin-specific Th17 cells, leading to enhanced disease severity and earlier onset. This exacerbated disease phenotype was previously associated with GM-CSF-mediated recruitment and activation of CD11b^+^Ly-6C^hi^ inflammatory monocytes to the CNS, and future studies will focus on these potential pathogenic mechanisms [26, 27]. In any case, it is clear from the current data, together with the recently described protective role of the respiratory Bacteroidetes species *P.melaninogenica* in a rat model of EAE [33], that common airway bacteria can regulate the development of extra-pulmonary disorders, particularly CNS autoimmune inflammation. Manipulation of the airway microbiome as an approach to dampen the pathogenicity of circulating myelin-specific T cells and therapeutic interventions that block T cell trafficking to the lungs should be considered crucial areas of MS research.

## Methods

### Animals

Wild-type C57BL/6J mice were bred and housed under SPF or germ-free (GF) conditions in the Comparative Medicine Unit at Trinity College Dublin. Sterility of GF mice was confirmed by routine microbiological screening of faecal samples and isolator swabs in aerobic and anaerobic conditions and qPCR amplification of the bacterial 16S rRNA gene using Eubacteria 16S rRNA (total bacteria) primers (UniF340: ACTCCTACGGGAGGCAGCAGT, UniR514: ATTACCGCGGCTGCTGGC) in fresh faecal samples. Bacterial genomic DNA was extracted using the QIAamp DNA Stool mini kit (Qiagen). Mice were 8-12 weeks old at initiation of experiments. All animal experiments were conducted under licence from the Health Products Regulatory Authority (HPRA) in Ireland and the University of Michigan Committee on Use and Care of Animals.

### Induction and Assessment of EAE

SPF mice were primed by s.c. injection at two sites on the back with 100μg of MOG_35-55_ (GenScript) emulsified 1:1 in 100μl CFA (Chondrex Inc.), containing 4mg/ml H37Ra Mtb. 10-12d post-immunisation, LN and spleen cells were were cultured at 10×10^6^/ml in RPMI containing β-mercaptoethanol (55μM), MOG (100ug/ml), IL-1β (10ng/ml), IL-23 (10ng/ml), αIL-4 (10ug/ml) and αIFNγ (10ug/ml) and cultured for 72h. Cells were enriched using CD4 magnetic beads (CD4 L3T4 Kit, Miltenyi Biotec) and 1X10^6^ CD4^+^ cells transferred i.p. to naïve mice. Animals were weighed daily and monitored for signs of clinical disease. Disease severity was graded as follows: 0-normal; 1-flaccid tail; 2-wobbly gait; 3-hind limb weakness; 4-paralysis of both hind limbs; 5-tetraparalysis/death. Only mice that developed clinical disease were included in data analysis.

### Cell isolation from tissues

For cell isolation from CNS tissues, mice were anaesthetized and perfused intracardially with 20ml ice-cold PBS and tissues isolated to HBSS containing 5% FBS before enzymatic digestion in Collagenase D (1mg/ml; Sigma) and DNase1 (100ng/ml; Applichem Lifesciences) and mononuclear cell isolation by percoll density centrifugation. Lungs were enzymatically digested in Collagenase D and DNase1, passed through a 70μm cell strainer and RBCs lysed with ACK lysis buffer (Invitrogen). LNs and Peyer’s patches were mechanically dissociated through a 70μm cell strainer using a syringe barrel. All cells were washed and counted before restimulation or flow cytometric analysis.

### Generation of murine bone marrow-derived dendritic cells and macrophages

BMDC were isolated as described by Lutz, Kukutsch *et al*., and re-plated 10d post-differentiation at 1X10^6^ cell/ml in RPMI supplemented with 10% FBS and 20ng/ml GM-CSF [34]. BMM were isolated as described by Weischenfeldt and Porse and re-plated 6d post-differentiation at 1X10^6^ cell/ml in DMEM supplemented with 10% FBS and 5% L929-conditioned medium [35].

### Preparation of bacterial inocula

Bacterial strains were streaked from frozen stocks on appropriate agar for the indicated times (Supp. Table 1) and inocula prepared in sterile PBS at the indicated CFU/ml by measuring OD600nm and confirmation by culture of serial dilutions. BMDC, BMM or AM were rested in antibiotic-free media for 3h before exposure to bacteria at MOI 0.1, 1, 10, 100, or PBS alone, and incubated at 37°C/5% CO_2_ for 24h. After 1h, cells were washed once with media containing gentamicin (100 μg/ml) and re-cultured in fresh media containing gentamicin for a further 23h. 24h post-exposure, supernatants were collected and stored at −20°C. For airway exposure of mice, 20μl of bacterial suspension in PBS (*V.parvula* or *M.catarrhalis* = 2X10^6^ CFU, *K.pneumoniae* = 1X10^6^ CFU) was administered i.n. to conscious naïve SPF or GF mice. Long-term monocolonisation with *K.pneumoniae* was initiated by adding 1X10^2^ CFU bacteria to drinking water in a sterile isolator. Colonisation and bacterial burden were regularly monitored in fecal samples and relevant tissues.

### Th17 cell co-culture with bacteria-stimulated BMDC

Following polarisation and CD4^+^ T cell enrichment, MOG-specific Th17 cells were rested for 72h before culture at 37°C/5% CO_2_ with BMDC (1:1) that had been exposed to the indicated bacteria at MOI 100 for 1h and washed in media containing gentamicin (100μg/ml). 24h later, supernatants were collected and stored at −20°C.

### ELISA

ELISA assays for IL-23, IL-12p70, TNFα IL-17A, IFNγ and GM-CSF (all R&D) were performed on cell culture supernatants, as per the manufacturer’s instructions. Cytotoxicity was determined by quantifying lactate dehydrogenase activity (CyQUANT LDH assay, ThermoFisher) in culture supernatants, relative to the maximal LDH activity of Triton-X100 lysed cells, according to the manufacturer’s instructions.

### Flow cytometry

Polarized Th17 cells or single cell suspensions isolated from lung, spleen and CNS tissue were activated (2×10^6^ cells/mL) with PMA (50ng/mL) and ionomycin (0.5μg/mL), in the presence of brefeldin A (5μg/mL; all Sigma), for 6h at 37°C/5% CO_2_. Activated cells, BMDC, BMM or AM, were washed and treated with Fcγ block (50 μg/mL) before extracellular staining with fluorochrome-conjugated antibodies against F4/80 (BM8), CD11b (M1/70), CD11c (N418), MHCII (M5/114.15.2), SiglecF (E50-2440), CD3 (17A2), CD4 (GK1.5), CD45.1 (A20) (Invitrogen or Biolegend). For intracellular cytokine staining, cells were fixed and permeabilized with the Fix&Perm Cell Permeabilisation Kit (Life Technologies) and stained with fluorochrome-conjugated antibodies against IL-17A (TC11-18H10.1), IFNγ (35F5) and GM-CSF (MP1-22E9) (Invitrogen or Biolegend). Viable cells were identified using LIVE/DEAD Aqua stain. Data was acquired on FACSCantoII or LSRFortessa flow cytometers (Becton Dickinson) and analyzed with FloJo software (Tree Star, Inc.), with gating set using fluorescence minus one (FMO) controls. Absolute numbers of donor and host cells were determined by multiplying the total viable cell count per tissue by the frequency of congenic cells among the total CD4^+^ T cell population, within the live cell gate.

### RT-PCR

Total RNA was extracted from tissues using the TRIzol/chloroform method, reverse transcribed into cDNA using a High-Capacity cDNA Reverse Transcription Kit (Applied Biosystems) and transcripts quantified by real-time semi-quantitative PCR using KiCqStart primers (Sigma) and iTaq SYBR green (Accuscience) on an iCycler PCR machine (Bio-Rad Laboratories). Data was normalized to the endogenous control 18S rRNA and expressed as fold change relative to controls.

### Statistics

Data was analysed for normal distribution using the Shapiro–Wilk test in GraphPad Prism(v9) software and statistical analysis between groups measured by unpaired two-tailed Student’s t test for parametric data or Mann–Whitney U test for nonparametric data. EAE clinical scores, mouse weight and donor cell accumulation in tissues over time were analysed by one- or two-way ANOVA with multiple comparisons. Data are shown as mean ± SEM. P-values of 0.05 or less were considered significant.

## Data Availability

The datasets generated during the current study are available from the corresponding author on reasonable request.

## Acknowledgements

This work was supported by research grants from Science Foundation Ireland (15/SIRG/3426) and the National Multiple Sclerosis Society (FG 1985-A-1) to SJL and a Wellcome Investigator Award (202846/Z/16/Z) to RMM. We thank Barry Moran for assistance with flow cytometry.

## Author Contributions

SJL conceived and supervised the study. All authors substantially contributed to the acquisition, analysis or interpretation of data and writing or critical review of the manuscript.

---

**Supplemental Figure 1.**
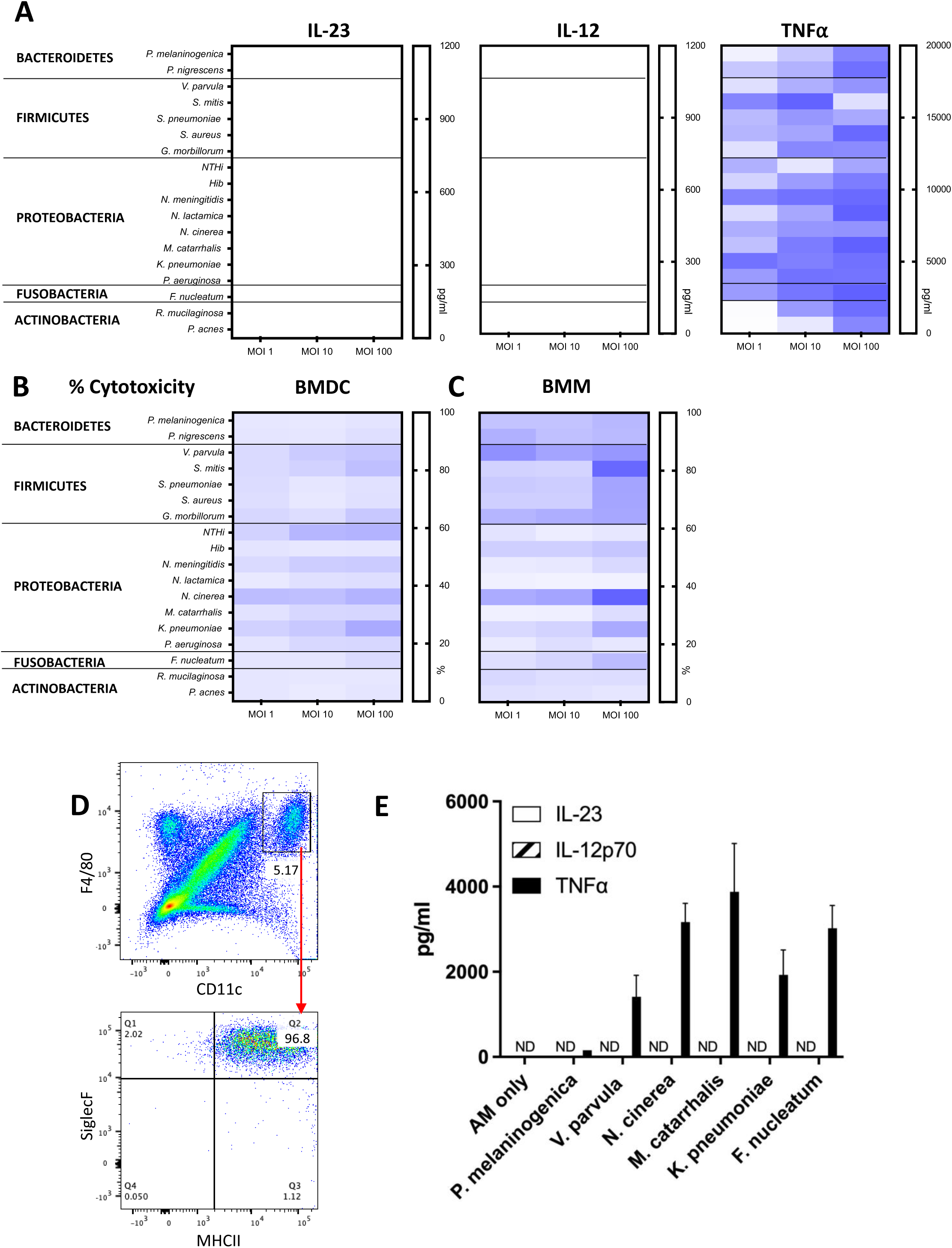
Selected respiratory tract bacteria do not induce IL-23 secretion by bone marrow derived macrophages *in vitro* or AM *ex vivo*. BMDC and BMM were seeded in triplicate at 2X10^5^ cells/well in a 96 well plate and rested for 3h before exposure to the indicated bacteria at MOI 1, 10 and 100. Cells were treated with gentamicin (100μg/ml) after 1h and supernatants collected 23h later. The concentration of IL-23, IL-12p70 and TNFα secretion by BMM was quantified by ELISA. Results are presented as a heat map indicating average cytokine concentration ml^−1^ (A; n = 3-9/group). Cytotoxicity was determined by quantifying lactate dehydrogenase (LDH) activity in BMDC (B) and BMM (C) culture supernatants and is displayed as percentage cytotoxicity of each species relative to the maximal LDH activity of Triton-X100 lysed cells. Lungs were harvested from naïve C57BL/6J mice and digested using Collagenase D and DNAse I. Cells were stained with sterile fluorochrome-conjugated antibodies against CD11b, CD11c, MHCII, F4/80 and Siglec F and sorted by flow cytometry. Representative FACS plots for one individual experiment are shown (D). AM were resuspended in cDMEM and seeded at a concentration of 5X10^4^ cells/well in a 96 well plate and rested for 3h. Cells were stimulated with selected bacteria at MOI 100 for 1h before treatment with Gentamicin (100 μg/ml), and supernatants collected 23h later. Concentrations of the cytokines IL-23, IL-12p70 and TNFα were quantified by ELISA. Results are expressed as mean ± SEM (E; n = 4-6/group).

**Supplemental Figure 2.**
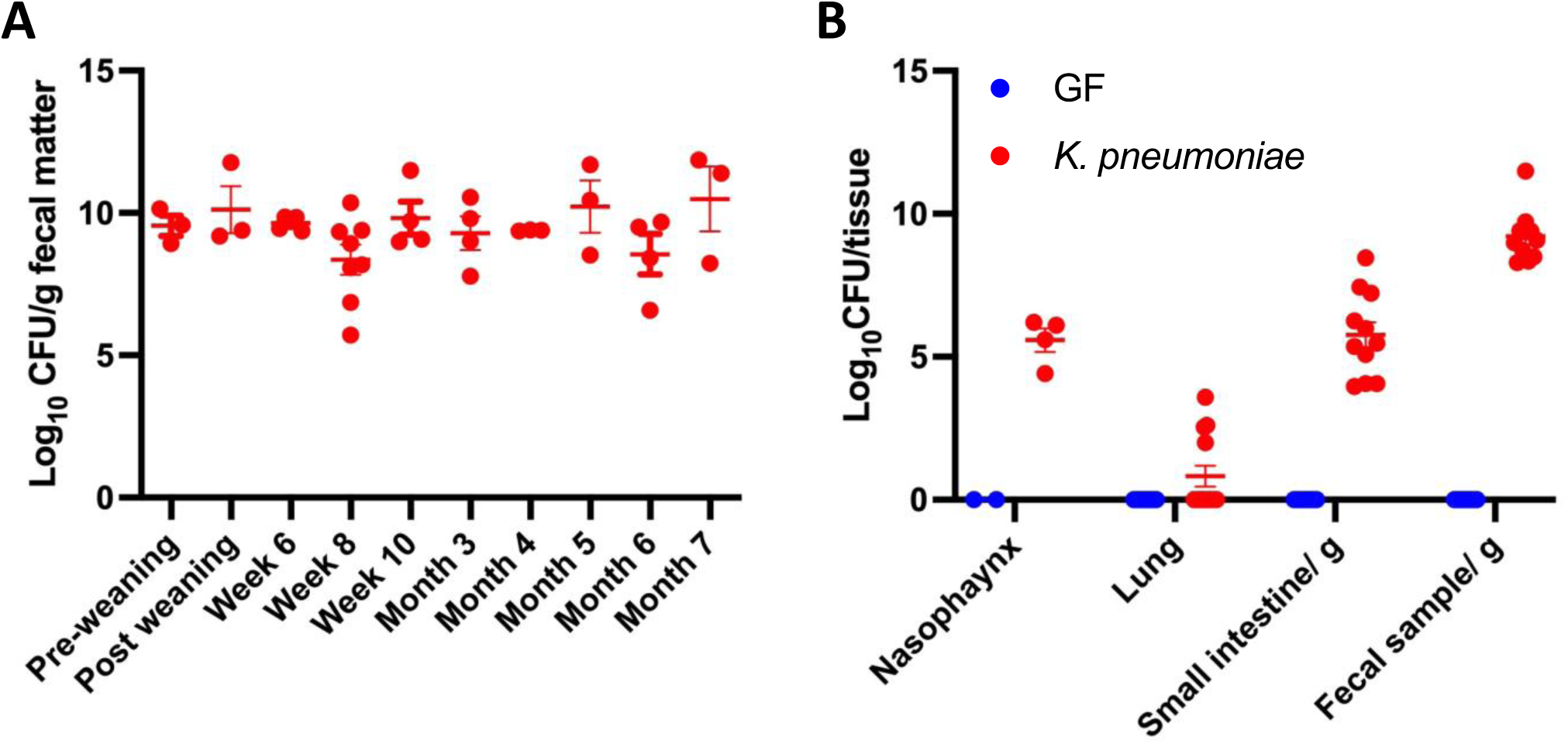
*K. pneumoniae* stably colonises the airways and gastrointestinal tract of gnotobiotic mice. *K. pneumoniae* was introduced to GF mice in the drinking water. Mice were then bred and maintained under monocolonised conditions in a gnotobiotic isolator for 7 months. Fresh faecal pellets were routinely collected, and lungs, nasopharyngeal tissue, small intestine and colon harvested from mice throughout life to assess bacterial burden. A) Data is expressed as Log_10_ CFU/g fecal matter. B) Data represents 8-week-old male and female mice and expressed as Log_10_ CFU/ tissue. Results are expressed as mean ± SEM.

**Supplemental Figure 3.**
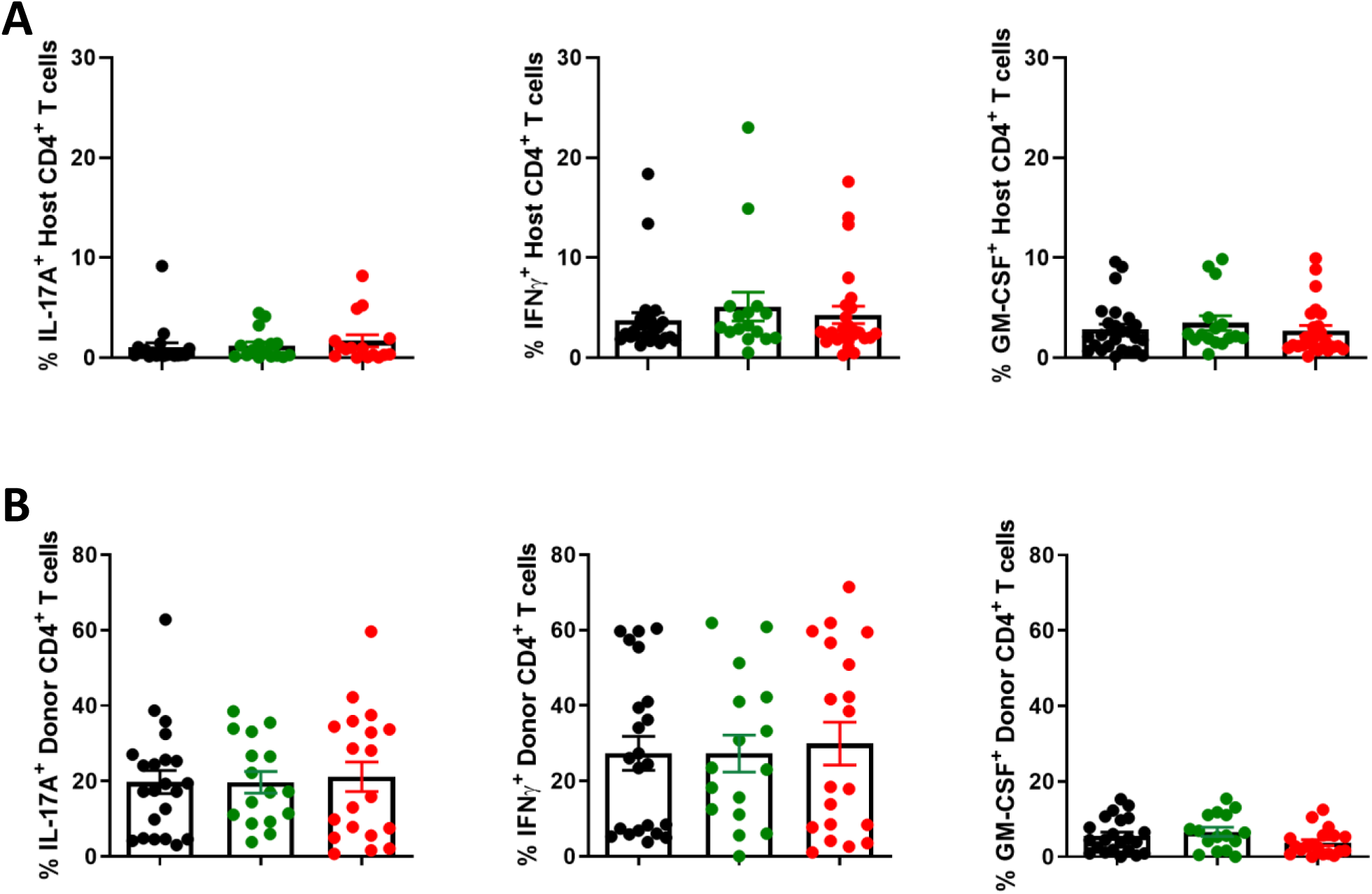
Exposure to *K.pneumoniae* and *M.catarrhalis* does not affect cytokine expression by host CD4 T cells in the lungs or donor CD4 T cells in the spleen of recipient mice. CD45.1^+^ donor Th17 cells were adoptively transferred (1X10^6^ CD4^+^ T cells/mouse) to congenic CD45.2 SPF hosts via i.p. injection. 2d after transfer, groups of T cell recipient mice were exposed to bacteria via i.n. administration of *M.catarrhalis* or *K.pnumoniae*, or PBS alone. 6d post-transfer (pre-clinical), lung and spleen tissues harvested. Single cell suspensions were stimulated with PMA and ionomycin with brefeldin A for 4 h, stained for surface CD3, CD4, CD45.1 and intracellular IL-17A, IFNγ and GM-CSF, and analysed by flow cytometry. Results are mean frequency of host IL-17A^+^, IFNγ^+^ and GM-CSF^+^ CD4 T cells in the lungs (A) or donor IL-17A^+^, IFNγ^+^ and GM-CSF^+^ CD4 T cells in the spleen (B) +/− SEM (n = 16-25/group from 4-5 independent experiments).

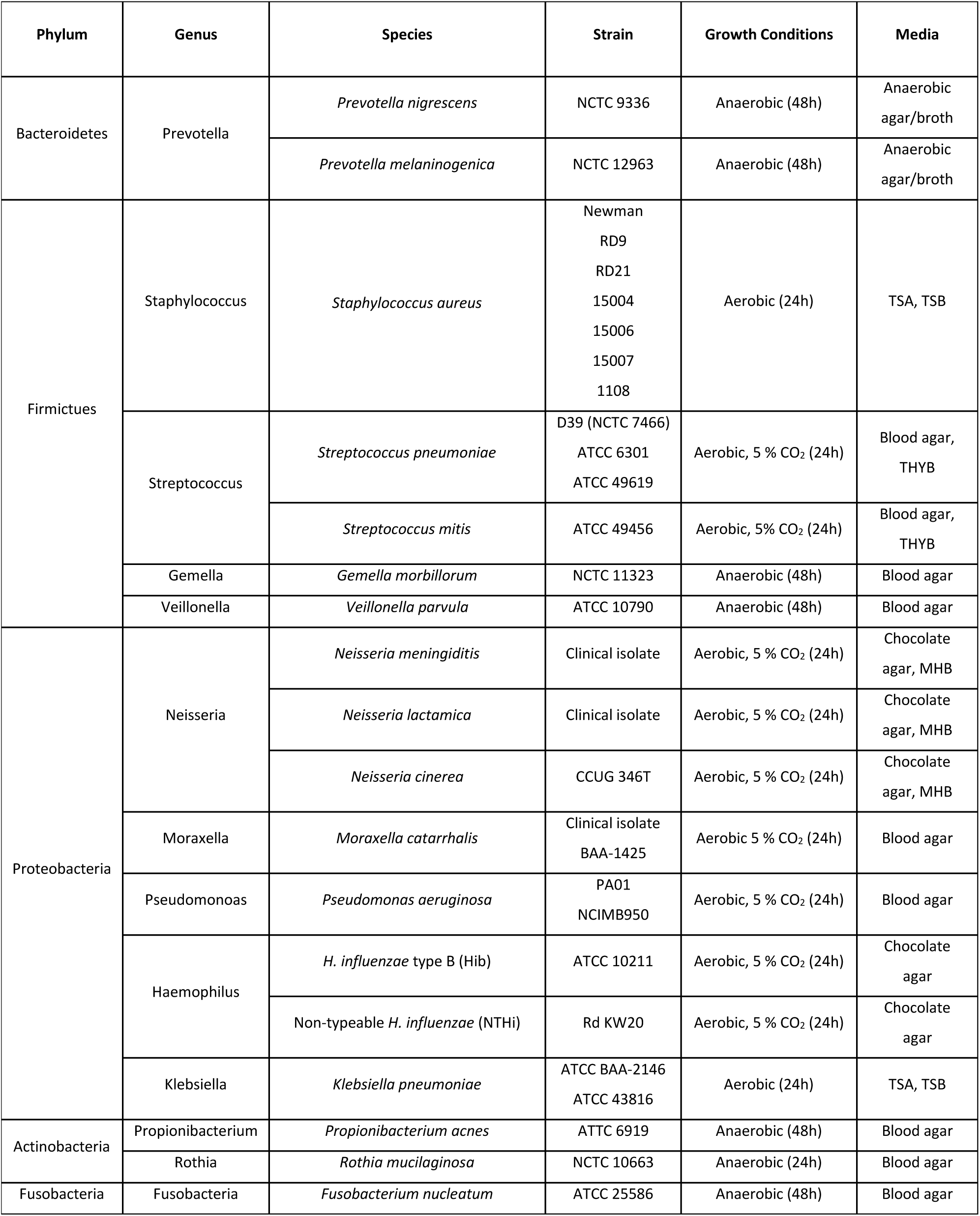
Supplemental Table 1

Supplemental Table 1
| Phylum | Genus | Species | Strain | Growth Conditions | Media |
| --- | --- | --- | --- | --- | --- |
| Bacteroidetes | Prevotella | *Prevotella nigrescens* | NCTC 9336 | Anaerobic (48h) | Anaerobic agar/broth |
|  |  | *Prevotella melaninogenica* | NCTC 12963 | Anaerobic (48h) | Anaerobic agar/broth |
| Firmicutes | Staphylococcus | *Staphylococcus aureus* | Newman<br>RD9<br>RD21<br>15004<br>15006<br>15007<br>1108 | Aerobic (24h) | TSA, TSB |
|  | Streptococcus | *Streptococcus pneumoniae* | D39 (NCTC 7466)<br>ATCC 6301<br>ATCC 49619 | Aerobic, 5 % CO<sub>2</sub> (24h) | Blood agar,<br>THYB |
|  |  | *Streptococcus mitis* | ATCC 49456 | Aerobic, 5% CO<sub>2</sub> (24h) | Blood agar,<br>THYB |
|  | Gemella | *Gemella morbillorum* | NCTC 11323 | Anaerobic (48h) | Blood agar |
|  | Veillonella | *Veillonella parvula* | ATCC 10790 | Anaerobic (48h) | Blood agar |
| Proteobacteria | Neisseria | *Neisseria meningitidis* | Clinical isolate | Aerobic, 5 % CO<sub>2</sub> (24h) | Chocolate agar, MHB |
|  |  | *Neisseria lactamica* | Clinical isolate | Aerobic, 5 % CO<sub>2</sub> (24h) | Chocolate agar, MHB |
|  |  | *Neisseria cinerea* | CCUG 346T | Aerobic, 5 % CO<sub>2</sub> (24h) | Chocolate agar, MHB |
|  | Moraxella | *Moraxella catarrhalis* | Clinical isolate<br>BAA-1425 | Aerobic 5 % CO<sub>2</sub> (24h) | Blood agar |
|  | Pseudomonas | *Pseudomonas aeruginosa* | PA01<br>NCIMB950 | Aerobic, 5 % CO<sub>2</sub> (24h) | Blood agar |
|  | Haemophilus | *H. influenzae* type B (Hib) | ATCC 10211 | Aerobic, 5 % CO<sub>2</sub> (24h) | Chocolate agar |
|  |  | Non-typeable *H. influenzae* (NTHi) | Rd KW20 | Aerobic, 5 % CO<sub>2</sub> (24h) | Chocolate agar |
|  | Klebsiella | *Klebsiella pneumoniae* | ATCC BAA-2146<br>ATCC 43816 | Aerobic (24h) | TSA, TSB |
| Actinobacteria | Propionibacterium | *Propionibacterium acnes* | ATCC 6919 | Anaerobic (48h) | Blood agar |
|  | Rothia | *Rothia mucilaginosa* | NCTC 10663 | Anaerobic (24h) | Blood agar |
| Fusobacteria | Fusobacteria | *Fusobacterium nucleatum* | ATCC 25586 | Anaerobic (48h) | Blood agar |

## References

1. Mannion JM., M.R., Lalor SJ, The Airway Microbiome-IL-17 Axis: a Critical Regulator of Chronic Inflammatory Disease. Clinical Reviews in Allergy and Autoimmunity, 2022, DOI: 10.1007/s12016-022-08928-y.

2. Hilty, M., et al., Disordered microbial communities in asthmatic airways. PLoS One, 2010. 5(1): p. e8578.

3. Erb-Downward, J.R., et al., Analysis of the lung microbiome in the “healthy” smoker and in COPD. PLoS One, 2011. 6(2): p. e16384.

4. Cox, M.J., et al., Airway microbiota and pathogen abundance in age-stratified cystic fibrosis patients. PLoS One, 2010. 5(6): p. e11044.

5. Demoruelle, M.K., et al., The Lung Microbiome Differs in Asymptomatic Subjects at Elevated Risk of Future Rheumatoid Arthritis Compared with Healthy Control Subjects. Annals of the American Thoracic Society, 2014. 11: p. S74.

6. Odoardi, F., et al., T cells become licensed in the lung to enter the central nervous system. Nature, 2012. 488(7413): p. 675–9.

7. Kanayama, M., Y.W. He, and M.L. Shinohara, The lung is protected from spontaneous inflammation by autophagy in myeloid cells. J Immunol, 2015. 194(11): p. 5465–71.

8. Edwards, S.C., S.C. Higgins, and K.H.G. Mills, Respiratory infection with a bacterial pathogen attenuates CNS autoimmunity through IL-10 induction. Brain, Behavior, and Immunity, 2015. 50: p. 41–46.

9. Glenn, J.D., C. Liu, and K.A. Whartenby, Frontline Science: Induction of experimental autoimmune encephalomyelitis mobilizes Th17-promoting myeloid derived suppressor cells to the lung. J Leukoc Biol, 2019. 105(5): p. 829–841.

10. McDonald, W.I. and T.A. Sears, The effects of experimental demyelination on conduction in the central nervous system. Brain, 1970. 93(3): p. 583–98.

11. Rao, P. and B.M. Segal, Experimental autoimmune encephalomyelitis. Methods Mol Biol, 2012. 900: p. 363–80.

12. Willer, C.J., et al., Twin concordance and sibling recurrence rates in multiple sclerosis. Proc Natl Acad Sci U S A, 2003. 100(22): p. 12877–82.

13. Ingelfinger, F., et al., Twin study reveals non-heritable immune perturbations in multiple sclerosis. Nature, 2022. 603(7899): p. 152–158.

14. Correale, J., M. Fiol, and W. Gilmore, The risk of relapses in multiple sclerosis during systemic infections. Neurology, 2006. 67(4): p. 652–9.

15. Berer, K., et al., Commensal microbiota and myelin autoantigen cooperate to trigger autoimmune demyelination. Nature, 2011. 479(7374): p. 538–41.

16. Lee, Y.K., et al., Proinflammatory T-cell responses to gut microbiota promote experimental autoimmune encephalomyelitis. Proc Natl Acad Sci U S A, 2011. 108 Suppl 1(Suppl 1): p. 4615–22.

17. Brucklacher-Waldert, V., et al., Phenotypical and functional characterization of T helper 17 cells in multiple sclerosis. Brain, 2009. 132(Pt 12): p. 3329–41.

18. Kebir, H., et al., Preferential recruitment of interferon-gamma-expressing TH17 cells in multiple sclerosis. Ann Neurol, 2009. 66(3): p. 390–402.

19. Huber, A.K., et al., Dysregulation of the IL-23/IL-17 axis and myeloid factors in secondary progressive MS. Neurology, 2014. 83(17): p. 1500–7.

20. Lock, C., et al., Gene-microarray analysis of multiple sclerosis lesions yields new targets validated in autoimmune encephalomyelitis. Nat Med, 2002. 8(5): p. 500–8.

21. Matusevicius, D., et al., Interleukin-17 mRNA expression in blood and CSF mononuclear cells is augmented in multiple sclerosis. Mult Scler, 1999. 5(2): p. 101–4.

22. McGeachy, M.J., et al., The interleukin 23 receptor is essential for the terminal differentiation of interleukin 17-producing effector T helper cells in vivo. Nat Immunol, 2009. 10(3): p. 314–24.

23. Hirota, K., et al., Fate mapping of IL-17-producing T cells in inflammatory responses. Nat Immunol, 2011. 12(3): p. 255–63.

24. Codarri, L., et al., RORγt drives production of the cytokine GM-CSF in helper T cells, which is essential for the effector phase of autoimmune neuroinflammation. Nat Immunol, 2011. 12(6): p. 560–7.

25. El-Behi, M., et al., The encephalitogenicity of T(H)17 cells is dependent on IL-1- and IL-23-induced production of the cytokine GM-CSF. Nat Immunol, 2011. 12(6): p. 568–75.

26. Amorim, A., et al., IFNγ and GM-CSF control complementary differentiation programs in the monocyte-to-phagocyte transition during neuroinflammation. Nature Immunology, 2022. 23(2): p. 217–228.

27. King, I.L., T.L. Dickendesher, and B.M. Segal, Circulating Ly-6C+ myeloid precursors migrate to the CNS and play a pathogenic role during autoimmune demyelinating disease. Blood, 2009. 113(14): p. 3190–7.

28. Duncker, P.C., et al., GM-CSF Promotes Chronic Disability in Experimental Autoimmune Encephalomyelitis by Altering the Composition of Central Nervous System-Infiltrating Cells, but Is Dispensable for Disease Induction. J Immunol, 2018. 200(3): p. 966–973.

29. McGeachy, M.J., et al., TGF-beta and IL-6 drive the production of IL-17 and IL-10 by T cells and restrain T(H)-17 cell-mediated pathology. Nat Immunol, 2007. 8(12): p. 1390–7.

30. Pollak, Y., et al., Behavioral aspects of experimental autoimmune encephalomyelitis. J Neuroimmunol, 2000. 104(1): p. 31–6.

31. Komuczki, J., et al., Fate-Mapping of GM-CSF Expression Identifies a Discrete Subset of Inflammation-Driving T Helper Cells Regulated by Cytokines IL-23 and IL-1β. Immunity, 2019. 50(5): p. 1289–1304.e6.

32. Caucheteux, S.M., et al., Cytokine regulation of lung Th17 response to airway immunization using LPS adjuvant. Mucosal Immunol, 2017. 10(2): p. 361–372.

33. Hosang, L., et al., The lung microbiome regulates brain autoimmunity. Nature, 2022.

34. Lutz, M.B., et al., An advanced culture method for generating large quantities of highly pure dendritic cells from mouse bone marrow. J Immunol Methods, 1999. 223(1): p. 77–92.

35. Weischenfeldt, J. and B. Porse, Bone Marrow-Derived Macrophages (BMM): Isolation and Applications. CSH Protoc, 2008. 2008: p. pdb.prot5080.

